# A global comparison of the nutritive values of forage plants grown in contrasting environments

**DOI:** 10.1101/224808

**Authors:** Mark A. Lee

## Abstract

Forage plants are valuable because they maintain wild and domesticated herbivores, and sustain the delivery of meat, milk and other commodities. Forage plants contain different quantities of fibre, lignin, minerals and protein, and vary in the proportion of their tissue that can be digested by herbivores. These nutritive components are important determinants of consumer growth rates, reproductive success and behaviour. A dataset was compiled to quantify variation in forage plant nutritive values within- and between-plant species, and to assess variation between plant functional groups and bioclimatic zones. 1,255 geo-located records containing 3,774 measurements of nutritive values for 136 forage plant species grown in 30 countries were obtained from published articles. Spatial variability in forage nutritive values indicated that climate modified plant nutritive values. Forage plants grown in arid and equatorial regions generally contained less digestible material than those grown in temperate and tundra regions; containing more fibre and lignin, and less protein. These patterns may reveal why herbivore body sizes, digestion and migration strategies are different in warmer and drier regions. This dataset also revealed the capacity for variation in the nutrition provided by forage plants. The proportion of the plant tissue that was digestible ranged between species from 2-91%. The amount of fibre contained within plant material ranged by 23-90%, protein by 2-36%, lignin by 1-21% and minerals by 2-22%. Water contents also varied substantially; ranging from 3-89% of standing biomass. On average, grasses and tree foliage contained the most fibre, whilst herbaceous legumes contained the most protein and tree foliage contained the most lignin. However, there were individual species within each functional group that were highly nutritious. This dataset may be used to identify forage plant species with useful traits which can be cultivated to enhance livestock productivity and inform wild herbivore conservation strategies.

## Introduction

Forage plants provide humans with valuable ecosystem services, for example, they feed an estimated 1.5 billion cattle, 1.2 billion sheep, 1 billion goats and 0.2 billion buffalo around the world – supplying meat, milk and other commodities (FAOSTAT 2016). These livestock are a global asset, worth around $1.4 trillion to the global economy, and livestock farming employs around 1.3 billion people, directly supporting over 600 million smallholder farmers (Thornton et al. 2011). Wild herbivores also feed on forage plants and therefore these plants contribute to the maintenance of biodiversity, to the complexity of biotic interactions and to the magnitude and direction of the associated ecosystem processes and services (Millennium Ecosystem Assessment 2005).

Plants vary in the quantities of different nutritive components that they deliver to consumers. They can vary in the amounts of fat, protein, carbohydrate, fibre and other micro-nutrients that are present in tissues. Herbivores vary in their requirements for these different nutritive components, and their dietary requirements change over time (Simpson et al. 2004). Forage plants also vary in their palatability, with defensive or structural compounds such as lignin and fibrous compounds reducing the amount of plant material that herbivores can digest (Distel et al. 2005). To reflect these different nutritive components, there are several agronomic metrics of forage nutritive quality. Metrics range from the quantification of forage dry matter content (DM: the proportion of plant material remaining after drying) to the assessment of forage digestibility (an integrative value estimating the proportion of plant material which can be digested by herbivores) (Gardarin et al. 2014; Beecher et al. 2015). Multiple nutritive metrics may be considered together to estimate the value of forage species or varieties to livestock and wild herbivores, and to project future milk or meat yields (Dong et al. 2003; Jégo et al. 2013). An understanding of the nutritive value of plants has been used to guide ecosystem management strategies, including forage species selection (Cherney and Cherney 1997; Delaby and Peyraud 2009).

Foraging theory links the diets of herbivores to their fitness, providing insights into patch selection, consumer population sizes and animal movements (Pyke 1984). Larger patch areas and enhanced plant biomass production have been positively correlated with consumer persistence, population sizes, and has been negatively correlated with rates of extinction (Hanski and Thomas 1994; Schlinkert et al. 2016). However, the quality and palatability of forage plants also affects the amount of vegetation that is consumed, rates of animal bodyweight gains and reproductive success (Herrero et al. 2015). The nutritive value of forage plants determines optimal herbivore body sizes, the relative success of ruminants and non-ruminants, and migration strategies (Bailey et al. 1996). The paucity of data quantifying the nutritive value of different forage plants grown across different locations means that nutrition is rarely considered as a part of ecological or conservation studies (Pontes et al. 2007). Plant species composition determines the nutritional quality of semi-natural grasslands (French 2017), alpine grasslands (Komac et al. 2014) and pasture (Chapman et al. 2014). Herbivores can consume herbaceous legumes and non-legumes, as well as the foliage of shrubs and trees (Wood et al. 2015). There is emerging evidence that there is variation in the nutritive values of plant functional groups. Herbaceous legumes may deliver greater quantities of protein and grasses may be more readily digestible (Weller and Cooper 2001; King et al. 2012). The extent by which forage plants from different functional groups can vary in their nutritive value and palatability has not been comprehensively assessed at the global scale. In a previous study focussing solely on grasses, Lee et al. (2017) demonstrated that the fibre and protein contents of forage grasses (55 species from 16 countries) ranged from 34-90% and from 5-36%, respectively. Incomplete data coverage means that comparisons between the nutritive values of forage plants grown in different regions have also not been fully quantified, although there is evidence that warmer regions are associated with lower quality forage grasses, containing higher proportions of fibre, which are generally tougher to digest (Lee et al. 2017).

To further extend data coverage, and to investigate the variation between functional groups and regions a new study was undertaken, and is presented here. Two main hypotheses were tested; firstly, that there would be considerable variation between species and functional groups, such as greater protein content in leguminous herbaceous plants and greater lignin content in the foliage of trees. The second hypothesis was that forage plants grown in hotter and drier regions would be of lower nutritive quality than those grown in cooler and wetter regions, containing higher proportions of fibre and lignin, lower proportions of protein and thus would be associated with lower digestibility values. To test these hypotheses, a large geo-referenced database of forage plants was compiled, which included a range of nutritive metrics. Nutritive metrics were compared within- and between-forage plant species and between functional groups and bioclimatic zones.

## Material and methods

### Nutritive metrics

The metrics that were chosen for inclusion in the database were the eight most commonly reported agronomic metrics in a pilot assessment of journal articles listed by the ISI Web of Knowledge (WoK; <u>www.wok.mimas.ac.uk</u>). For consistency, values were included in the database if they were derived from laboratory analyses and based on the methods of Van Soest et al. (1991) or AOAC (2000). Mineral ash values represented the mineral component of the forage plants (hereafter termed ‘ash’: the inorganic mineral component remaining following burning). Two fibre metrics were included, representing: (1) the plants structural components termed acid detergent fibre (ADF: the material remaining after boiling in acid detergent, representing lignin, cellulose, silica and insoluble nitrogenous compounds but not hemicellulose); and (2) termed neutral detergent fibre (NDF: the material remaining after boiling in neutral detergent representing lignin, silica, cellulose and hemicellulose). Lignin was included when it was presented as acid detergent lignin (ADL: isolated by boiling in strong acid). Forage protein content was included in the dataset when presented as crude protein (CP: total nitrogen content as measured by Kjeldahl digestion multiplied by 6.25). The dry matter contents of the forage plants was also included (DM: the proportion of material remaining following drying).

Two digestibility metrics were also included in the dataset, as integrated metrics estimating the proportion of forage that can be utilised by ruminants. Dry matter digestibility (DMD: the proportion of forage dry matter which can be digested) and organic matter digestibility (OMD: the proportion of forage organic matter which can be digested). Digestibility metrics were estimated using *in vitro*, *in vivo* and near infra-red (NIR) techniques.

### Data collection

Data were obtained from peer-reviewed journal articles. These articles were identified by systematically searching the WoK. To avoid researcher bias and to maintain a consistent approach, the search terms used to identify the articles listed in the WoK were identified a priori. Articles were included in the database if the nutritive measurements were related to a specific forage plant species or hybrid that had been grown in field conditions at a defined location (hereafter termed ‘site’) and harvested for nutritional analyses at a stated time. Data from experiments conducted in greenhouses or field experiments, i.e. those that manipulated climatic variables, were excluded because the prevailing growing conditions were not representative of the location. All plant species names were checked for accuracy using an online list of species names, with synonyms switched to accepted names and unknown species were removed (<u>www.theplantlist.org</u>).

To ensure that the methods for measuring forage nutritive value were consistent across the articles, data were included if Ash, ADF, ADL, DM, NDF and/or CP analyses were carried out on dried samples and presented in units of g kg^−1^ DM or % DM. DMD and OMD was also recorded when available. All measurements that were taken at the same site and on the same sampling interval were allocated to the same row of the dataset, thus multiple nutritive metrics were included for the same time and location (mean nutritive metrics per row = 3.01 ± 0.04). Samples were included if they were analysed in the same form as they would be consumed by livestock; grasses, herbaceous non-legumes (hereafter termed ‘herbs’) and herbaceous legumes (hereafter termed ‘legumes’) were included as whole plants, whilst trees and shrubs were included if analyses were carried out on foliage. For our analyses, the foliage of trees and shrubs were grouped together (hereafter termed ‘tree’).

Sites were allocated to a bioclimatic zone as defined by the Köppen–Geiger climate classification system (Kottek et al. 2006) and recorded in the database as arid (≥70 % of precipitation falls in summer or winter), equatorial (mean temperature of the coldest month ≥18 °C), temperate (mean temperature of the warmest month ≥ 10 °C and the coldest month −3−18 °C) or tundra (mean temperature of the warmest month ≥ 10 °C and the coldest month ≤ −3 °C). Hot and dry zones (arid and equatorial) and cool and wet zones (temperate and tundra) were grouped together (for details of the sites included in the database see Supplementary Material 1).

### Representation in the database

The database contained 1,255 geo-located records with 3,774 measurements of nutritive values for 136 forage plant species or hybrid cultivars grown in 30 countries (for a summary of all of the mean nutritive values across all plant species see Supplementary Material 2). The most commonly recorded nutritive metric was CP, which was measured in 88% of the records and in all 30 countries. This was followed by the two fibre metrics, ADF and NDF, which were measured in 65% and 64% of the records (22 and 25 of the countries), respectively. ADL, Ash and DM were less commonly recorded, and were present in 20%, 16% and 20% of the records (13, 15 and 14 countries), respectively. Of the two digestibility metrics, DMD was recorded more than twice as frequently as OMD, and they were both recorded from 14 and 9 countries, respectively.

Grasses were the most commonly recorded functional group, representing 87% of all records, with legumes, trees and herbs making up 10%, 3% and 1% of the dataset, respectively. Records were the most numerous from the tundra bioclimatic zone, comprising 49% of the dataset, compared with 33% from the temperate zone, 15% from the arid zone and 3% from the equatorial zone. However, temperate records were more likely to contain multiple nutritive metrics and therefore the temperate zone contributed the largest total number of measurements to the dataset (2035 values), followed by tundra (981 values), arid (541 values) and equatorial zones (217 values).

### Statistics

Nutritive metrics, Ash, ADF, ADL, DM, NDF and CP were correlated with both DMD and OMD using linear regression analyses, with degrees of fit for regression lines calculated using r^2^. In all cases either DMD or OMD was the response variable with the other metrics included as potential explanatory variables. Prior to statistical testing, data were tested for non-linearity by comparing quadratic and logarithmic models with linear models. In all cases linear models were the most appropriate. Variation between functional groups and bioclimatic zones for each nutritive metric was assessed using Analysis of Variance (ANOVA) tests, with significant differences between individual zones and groups identified using Tukey’s Honest Significant Different (HSD) tests. All analyses were computed using R version 3.2.3 (The R Foundation for Statistical Computing, Vienna, Austria, 2016).

## Results

### Comparisons of nutritive metrics

The mean DM across all of the forage plants was 41% and the mean water content of the plants was 59% (Table 1). In terms of the fibre content across the whole dataset, means values for ADF and NDF were 32% and 57%, respectively. Mean CP was the next highest value at 15%, with mean ash at 9% and mean ADL at 6%. Overall, of the plant material that was measured, a mean of 71% in terms of DMD, and a mean of 62% in terms of OMD, was digestible.

**Table 1:**
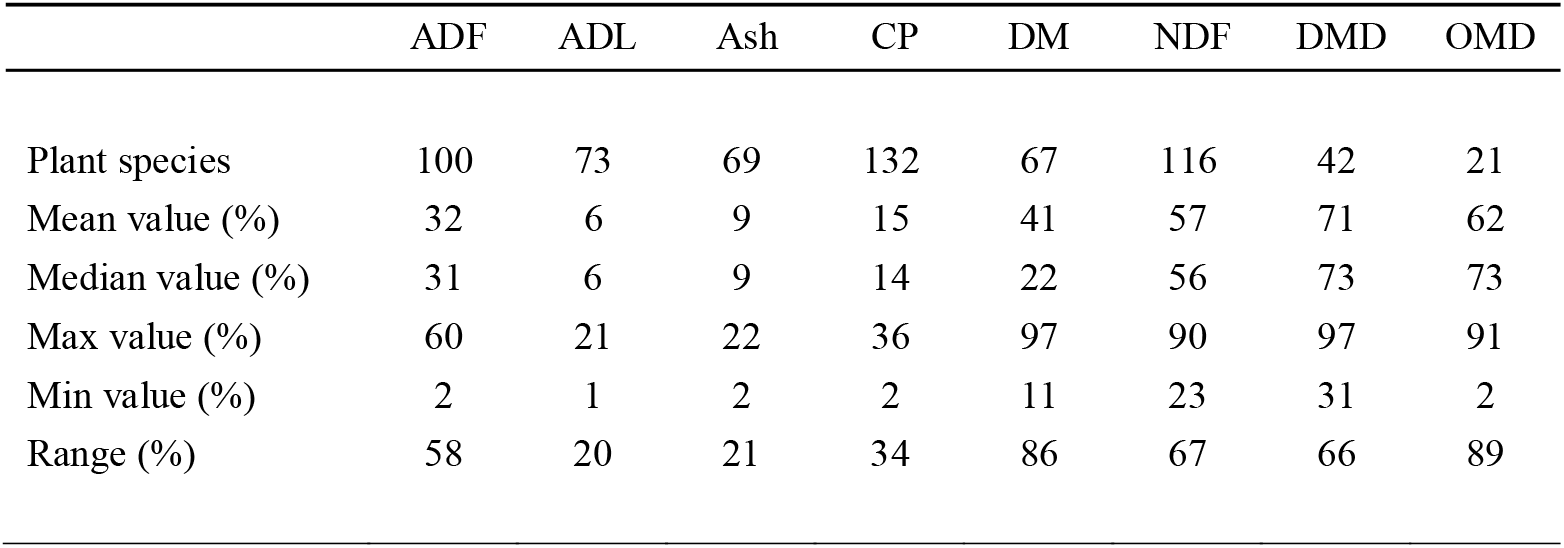
A count of the number of forage plant species in the database and the mean, median, maximum (max), minimum (min) and range of values across all of the records. Metrics are acid detergent fibre (ADF), acid detergent lignin (ADL), mineral ash (Ash), crude protein (CP), dry matter (DM), neutral detergent fibre (NDF), dry matter digestibility (DMD) and organic matter digestibility (OMD).

|  | ADF | ADL | Ash | CP | DM | NDF | DMD | OMD |
| --- | --- | --- | --- | --- | --- | --- | --- | --- |
| Plant species | 100 | 73 | 69 | 132 | 67 | 116 | 42 | 21 |
| Mean value (%) | 32 | 6 | 9 | 15 | 41 | 57 | 71 | 62 |
| Median value (%) | 31 | 6 | 9 | 14 | 22 | 56 | 73 | 73 |
| Max value (%) | 60 | 21 | 22 | 36 | 97 | 90 | 97 | 91 |
| Min value (%) | 2 | 1 | 2 | 2 | 11 | 23 | 31 | 2 |
| Range (%) | 58 | 20 | 21 | 34 | 86 | 67 | 66 | 89 |

There was a larger range of values for OMD than for DMD, with digestibility ranging from 2-91% and from 31-97%, for the two metrics, respectively. In terms of the other nutritive metrics, DM had the largest range of values, ranging from 11-97%, followed by NDF at 23-90%, ADF at 13-60% and CP at 2-36%. The metrics with the largest ranges also represented the largest number of different plant species, since CP was recorded from 132 species, NDF was recorded from 116 species and ADF was recorded from 100 species. The exception to this was DM which was recorded from 67 forage plant species.

Several of the nutritive metrics were correlated with DMD and OMD, but there were differences in the degree of fit around the regression lines and the direction of the relationships (Table 2). NDF was strongly negatively correlated with both DMD and OMD, as indicated by high r^2^ values. CP was the only metric which was positively correlated with digestibility, both in terms of DMD and OMD, though the degree of fit of the regression line for CP and OMD was relatively low. ADF was also negatively correlated with DMD and OMD but the amount of variation explained by the regression line, and thus the degree of fit, was much lower than for NDF. ADL and DM were also negatively correlated with OMD but the degree of fit was lower between DMD and these two metrics.

**Table 2:** Regression outputs of the relationships between dry matter digestibility (DMD) or organic matter digestibility (OMD) and acid detergent fibre (ADF), acid detergent lignin (ADL), mineral ash (ash), crude protein (CP), dry matter (DM) and neutral detergent fibre (NDF).

| Metric | Equation | t | DF | P | r<sup>2</sup> |
| --- | --- | --- | --- | --- | --- |
| ADF | DMD = -0.12 + 109 | -17.3 | 105 | < 0.001 | 0.74 |
| ADL | DMD = -0.15 + 70 | -4.9 | 73 | < 0.001 | 0.24 |
| Ash | DMD = -0.03 + 73 | -0.7 | 57 | 0.49 | 0.01 |
| CP | DMD = 0.18 + 40 | 15.9 | 153 | < 0.001 | 0.62 |
| DM | DMD = -0.09 + 87 | -5.2 | 71 | < 0.001 | 0.26 |
| NDF | DMD = -0.10 + 130 | -17.7 | 146 | < 0.001 | 0.68 |
| ADF | OMD = -0.12 + 91 | -4.8 | 67 | < 0.001 | 0.26 |
| ADL | OMD = -0.57 + 96 | -17.3 | 31 | < 0.001 | 0.90 |
| Ash | OMD = -0.34 + 108 | -2.8 | 12 | < 0.05 | 0.35 |
| CP | OMD = 0.12 + 43 | 2.6 | 81 | < 0.05 | 0.06 |
| DM | OMD = -0.04 + 79 | -4.7 | 30 | < 0.001 | 0.41 |
| NDF | OMD = -0.09 + 120 | -15.3 | 79 | < 0.001 | 0.75 |

### Geographical variation between functional groups

Fibre values of the forage plants grown in arid and equatorial regions were a mean of 18% and 11% higher than those grown in temperate and tundra region, as defined by NDF (Figure 1a) and ADF (Figure 1b), respectively. However, CP values of forage plants grown across these drier regions were a mean of 2% lower than for plants grown in temperate or tundra regions (Figure 1c). Forage plants in arid and equatorial regions also contained greater amounts of ADL; a mean 3% greater than temperate and tundra regions (Figure 1d). DM contents were generally higher (and thus water contents lower) and mineral ash content lower in arid and equatorial regions (Table 3). Both of the digestibility metrics were lower for plants grown in arid and equatorial regions; a mean of 77% and 78% of the plant material grown in temperate and tundra regions was digestible when compared with 30% and 54% of the plants grown in arid and equatorial regions, considering both DMD and OMD, respectively.

**Figure 1:**
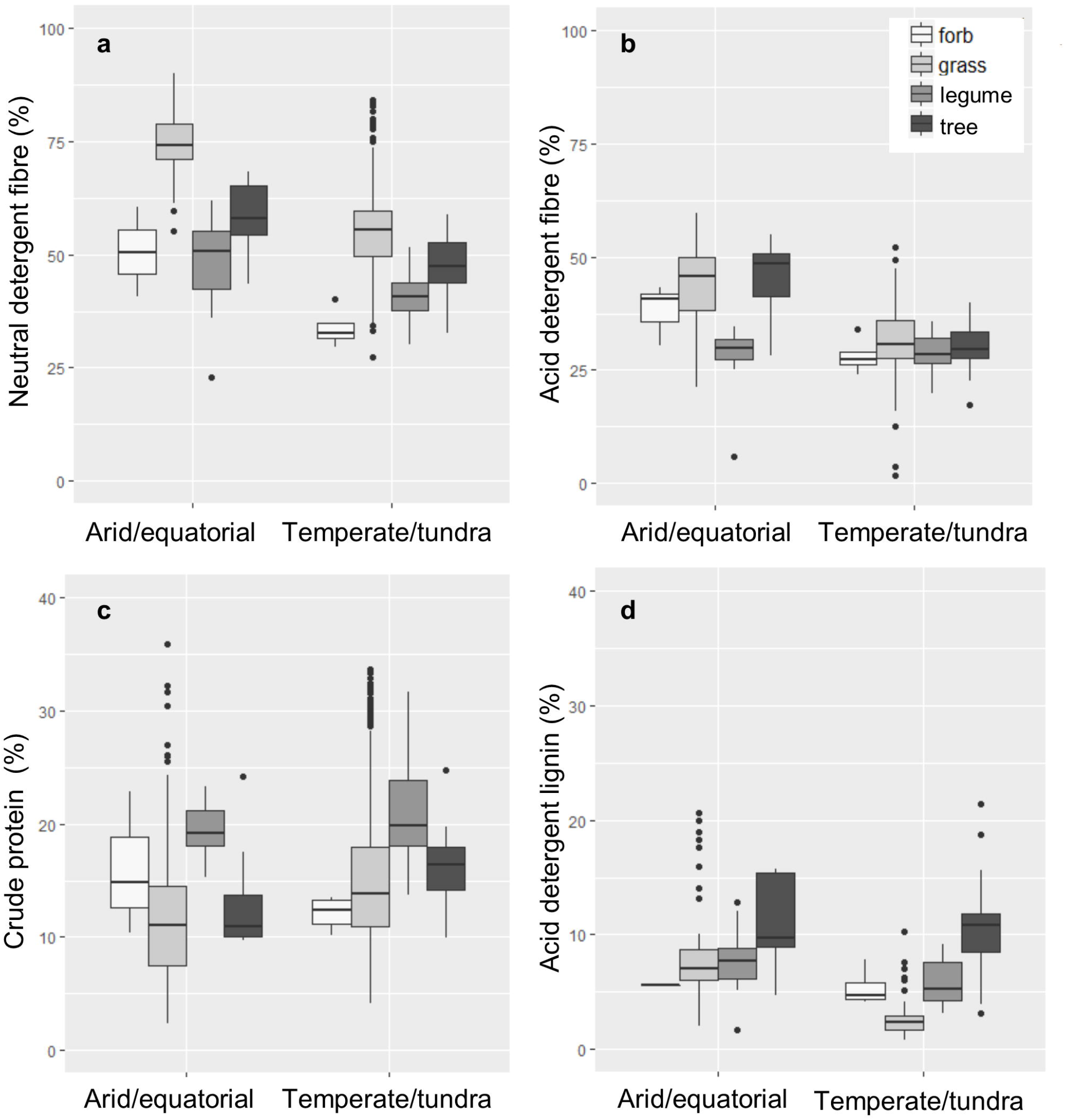
Boxplots representing the nutritive values of forage plants grown in arid and equatorial regions or temperature and tundra regions. Nutritive values are separated into plant functional groups; herbaceous non-legumes (herb), grasses, herbaceous legumes and trees. Metrics are (a) neutral detergent fibre, (b) acid detergent fibre, (c) crude protein and (d) acid detergent lignin.

**Table 3:** Pairwise comparisons of temperate and tundra (Te) and arid and equatorial (Eq) bioclimatic zones and pairwise comparisons of the functional groups; grass, herb, legume (leg) and tree, for each of the eight nutritive metrics. Positive values indicate that the first stated parameter in the pair is greater than the second, with associated P value.

|  | ADF |  | ADL |  | Ash |  | CP |  | NDF |  | DMD |  | OMD |  |
| --- | --- | --- | --- | --- | --- | --- | --- | --- | --- | --- | --- | --- | --- | --- |
|  | V | P | V | P | V | P | V | P | V | P | V | P | V | P |
| Te-Eq | -11 | <0.001 | -3 | <0.001 | 2 | <0.001 | 3 | <0.001 | -19 | <0.001 | 12 | <0.001 | 49 | <0.001 |
| grass-herb | 4 | >0.05 | -1 | >0.05 | -2 | >0.05 | 0 | >0.05 | 22 | <0.001 | -8 | >0.05 | - | - |
| leg-herb | 0 | >0.05 | 1 | >0.05 | 0 | >0.05 | 6 | <0.05 | 6 | >0.05 | 7 | >0.05 | - | - |
| tree-herb | 3 | >0.05 | 5 | <0.01 | -2 | >0.05 | 1 | >0.05 | 13 | <0.01 | 8 | >0.05 | - | - |
| leg-grass | -4 | <0.001 | 2 | >0.05 | 3 | <0.001 | 6 | <0.001 | -16 | <0.001 | 15 | >0.05 | -8 | >0.05 |
| tree-grass | 0 | >0.05 | 6 | <0.001 | 0 | >0.05 | 1 | >0.05 | -9 | <0.001 | 16 | >0.05 | -13 | >0.05 |
| tree-leg | 4 | <0.05 | 5 | <0.001 | -2 | <0.001 | -5 | <0.001 | 7 | <0.01 | 1 | >0.05 | -6 | >0.05 |

Grasses and tree foliage generally contained the most fibre; mean NDF was highest across the grasses at 59% and tree foliage at 50%, whilst NDF for legumes was the lowest with a mean of 42% (Figure 1a). Mean ADF displayed a similar pattern to NDF, with tree foliage having a mean ADF of 34% and the grasses having a mean of 33% (Figure 1b). As with NDF, legumes were the lowest in terms of ADF with a mean of 28%. Herbs were not significantly different from grasses, legumes or tree foliage in terms of either ADF or NDF.

Mean CP values for herbs, grasses and tree foliage were 14%, 15% and 15%, respectively – and were not significantly different from each other (Figure 1c). However, the mean CP value of legumes was greater than the other groups at 21%. The mean ADL value for tree foliage was between 5% and 6% greater than the other three functional groups (Figure 1d). Mean ash values of legumes were 2-3% greater than the grasses and tree foliage but not different from the herbs. There were no detectable differences in the digestibility of the functional groups, either in terms of DMD or OMD (Table 3).

## Capacity for variation within- and between-species

### Dry matter content

The DM content of the forage plants was highly variable. At the upper end of the range of values the grasses, *Cynodon nlemfuensis* and *Chloris pycnothrix* were both measured at 97% whilst *Cenchrus ciliaris* was measured at 96%. The foliage of three tree species were also very high in terms of DM, with *Grewia mollis, Capparis tomentosa* and *Leucaena leucocephala* all recorded at 93%. At the lower end of the scale, the lowest values were recorded from *Lolium perenne, Trifolium pratense* and *Medicago sativa* at 11%, 11% and 13%, respectively. The largest ranges of DM values that were recorded were from the grass, *Panicum maximum* (22-91%), the tree, *Leucaena leucocephala* (24-93%), the herbaceous legume, *Lablab purpureus* (43-91%) and the grass, *Lolium perenne* (11-37%).

### Fibre

There was also substantial variation in NDF values both within- and between-species (Figure 2). The largest absolute NDF values were recorded from the grasses; *Bouteloua gracilis* at 90%, *Aristida longiseta* at 88% and *Setaria macrostachya* at 86%. The maximum value recorded from any other functional group related to the foliage of two trees; *Bauhinia cheilantha* at 68% and *Mimosa caesapiniifolia* at 68%. NDF for tree foliage, herbs and legumes were clustered at the lower end of the range of values. The minimum values of NDF were recorded from the herbaceous legume, *Psophocarpus scandens* at 23%, the grass, *Dactylis glomerata* at 27% and the herb, *Sanguisorba minor* at 30%. The largest ranges of NDF values that were recorded were from the grasses; *Dactylis glomerata* (27-71%), *Phleum pratense* (36-68%), *Alopecurus pratensis* (39-70%) and *Lolium perenne* (34-62%).

**Figure 2:**
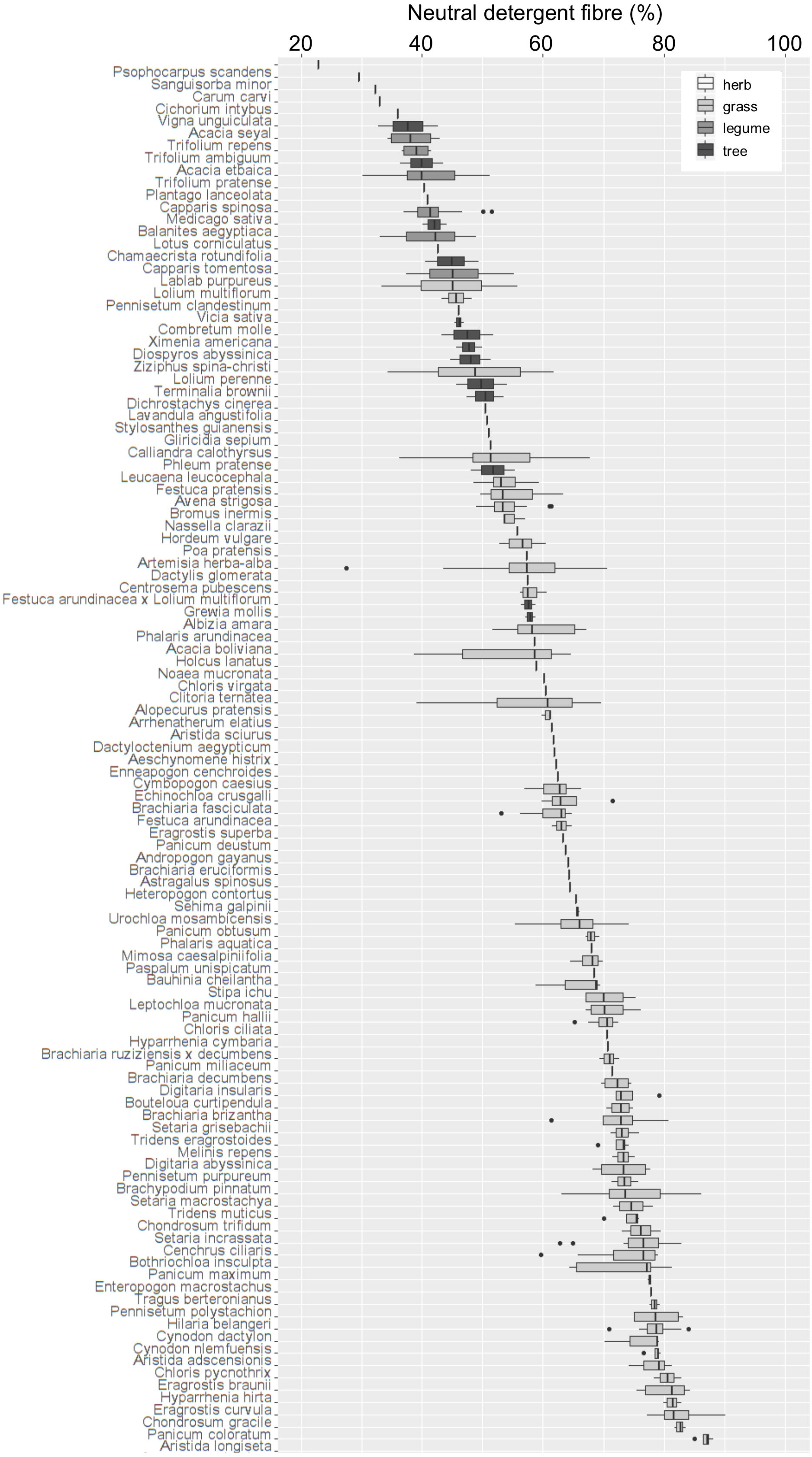
Ascending median neutral detergent fibre content for 116 forage plant species. Box shading represents functional group. Values are percent of dry plant material (% DM).

The largest ADF values were also measured from the grasses; *Hyparrhenia hirta* at 60% and *Enteropogon macrostachus* at 57%, whilst the foliage of the tree, *Mimosa caesapiniifolia*, was also recorded at 55%. High ADF values were rarer than high NDF and only 3% of ADF values in the database were greater than 50%. The lowest ADF values were measured from the grasses, *Phleum pratense, Agropyron riparium, Dactylis glomerata, Festuca arundinacea* and *Lolium multiflorum*, with values of 13%, 16%, 16%, 16% and 16%, respectively. The largest ranges of values were also measured from grasses; *Lolium perenne* (4-42%), *Lolium multiflorum* (2-35%), *Bromus inermis* (18-46%), *Dactylis glomerata* (16-44%) and *Phleum pratense* (13-38%).

### Protein

There was less variation in CP values compared with ADF and NDF values, both within- and between-species (Figure 3). The largest CP values were recorded from the grasses, *Agropyron cristatum* at 36% and *Lolium perenne* at 34%, the legume, *Medicago sativa* at 32%, the grass, *Elytrigia intermediate* at 32% and the herbaceous legume, *Trifolium repens* at 32%. The lowest CP values were recorded from the grasses, *Aristida adscensionis, Hyparrhenia hirta* and *Chloris-pycnothrix*; all at 2%. CP values for tree foliage, herbs and legumes were less clustered than for NDF but were more abundant towards the upper end of the range of values. The largest ranges of CP values were recorded from the grasses; *Agropyron cristatum* (8-36%), *Lolium perenne* (6-34%), *Lolium multiflorum* (6-28%) and *Elymus sibiricus* (5-26%).

**Figure 3:**
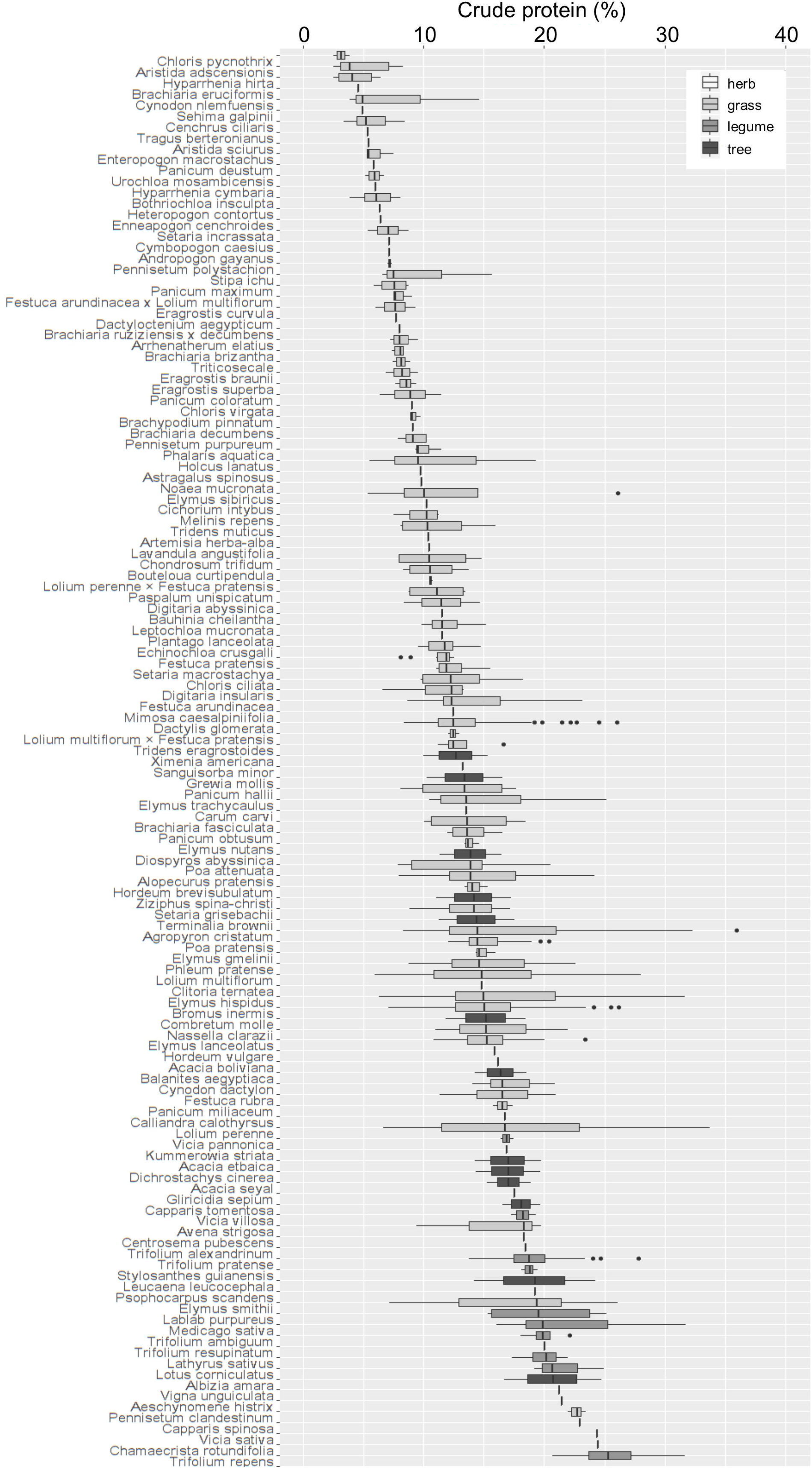
Ascending median crude protein content for 132 forage plant species. Box shading represents functional group. Values are percent of dry plant material (% DM).

### Mineral ash

The largest ash values were recorded across different functional groups, with maximum values recorded from the foliage of the tree, *Diospyros abyssinica* at 22%, the grass, *Pennisetum purpureum* at 19%, and the herbaceous legume, *Macroptilium atropurpureum* at 17%. High values were rare and only 4% of ash values were greater than 15%. Conversely, 70% of ash values were less than 10%, with minimum values of 2%, 2%, 2% and 4% recorded from the grasses, *Pennisetum purpureum*, *Pennisetum maximum* and *Brachiaria brizantha*, and from the foliage of *Bauhinia cheilantha*, respectively. The maximum ranges of ash values were recorded from the grasses, *Pennisetum purpureum* (2-18%), *Panicum maximum* (7-16%) and *Avena strigosa* (5-13%), as well as the foliage of two trees; *Terminalia brownie* (8-14%) and *Diospyros abyssinica* (16-22%).

### Lignin

The largest ADL values (i.e. those above 10%) were uncommon and represented only 12% of the dataset. Tree foliage of *Albizia amara* registered the greatest ADL content at 21%, with the grass *Brachiaria brizantha* and hybrid grass *Brachiaria ruziziensis x decumbens* having maximum ADL values of 21% and 19%, respectively. Foliage from *Grewia mollis* also had high ADL, with a maximum value of 19%. Low ash values were more common than high values across the dataset, with minimum values of 1% all recorded from the grasses; *Lolium multiflorum, Lolium perenne, Phleum pratense* and *Festuca arundinacea*, respectively. The ranges of values was also low for ADL, with the maximum ranges measured from the foliage of the tree *Albizia amara* (11-21%), the grass, *Setaria incrassate* (3-10%), the tree, *Grewia mollis* (12-19%) and the grass, *Chloris ciliata* (2-8%).

### Digestibility

The greatest absolute DMD values were recorded from the grass, *Phleum pratense* at 97%, with another grass, *Dactylis glomerata* at 90%, as well as the legumes, *Trifolium repens* at 89% and *Trifolium ambiguum* at 88%, also producing very high values. The largest DMD value for tree foliage was 79% for *Manihot pseudoglaziovii* and for the herbs it was 74% for *Carum carvi*. DMD values were recorded as low as 31% for *Hyparrhenia hirta*, 34% for *Aristida adscensionis*, 34% for *Enteropogon macrostachys* and 35% for *Enteropogon macrostachys* – all of which are grasses. The greatest ranges of DMD values were recorded for the grasses; *Elymus sibiricus* (47-85%), *Phleum pratense* (61-97%), *Hyparrhenia hirta* (31-64%) and *Lolium perenne* (56-86%).

There was a greater range of OMD values than DMD values, with the maximum OMD value recorded from *Lolium perenne* at 91%, with high values also recorded from the foliage of *Leucaena leucocephala* at 88%, the grasses, *Dactylis glomerata* and *Arrhenatherum elatius* each at 78%, with the hybrid grass *Festuca arundinacea x Lolium multiflorum* also reporting a high value of 77%. Low values of 3%, 4% and 9% were recorded from *Agropyron cristatum, Bromus inermis* and *Poa attenuata*, respectively. The largest ranges of OMD values were also recorded for the grasses; *Elymus sibiricus* (12-60%), *Lolium perenne* (61-91%), *Bromus inermis* (4-27%) and *Poa attenuata* (9-25%).

## Discussion

Forage plant nutrition is an important determinant of wild and domesticated herbivore population dynamics, plant/herbivore interactions and animal behaviour (Humphreys et al, 2005). Larger patch areas and enhanced plant biomass production have been correlated with larger and more persistent herbivore populations (e.g. Hanski & Thomas, 1994; Schlinkert *et al.*, 2016). However, if the currency of foraging theory is the provision of nutrition, then this dataset clearly demonstrates that individual plants or patches of plants with the same standing or dry biomass may be vastly different in terms of their nutritive values. This dataset shows that as much as 89% or as little as 3% of the standing biomass of forage plants is made up of water which dictates the amount of water which must be obtained from other water sources by consumers. These data also demonstrate that 91% of a forage plant may be digestible, compared with 2% for the least digestible plants (defined by OMD). Fibre (defined by NDF) can range by 23-90%, protein by 2-36%, lignin by 8-21% and minerals (defined by ash) by 2-22%. Such large variation in the nutritive values of forage plants changes the energetic costs of consumption versus the benefits of nutrient extraction for consumers.

Warmer regions have been associated with taller, less nutritious and slow-growing grasses (Jégo et al. 2013). Across all of the functional groups, this analysis showed that forage plants grown in warmer and drier regions were generally of lower nutritive value, as indicated by higher fibre, higher lignin and lower protein contents. These plants were also generally less readily digestible than those grown in cooler and wetter regions, as had been hypothesised. The reduced nutritive value of forage plants grown across these regions may be driven by increased abundances of plants with adaptations to avoid water loss and prevent heat stress. Adaptations include greater stem:leaf ratios, greater hair densities, thicker cell walls, more narrowly spaced veins, a higher proportion of epidermis, bundle sheaths, sclerenchyma and vascular tissues, and greater concentrations of lignin and silica (Kering et al. 2011).

Many species of arthropods, birds and mammals actively select or avoid plants based on their nutritive values (Greenberg and Bichier 2005; Amato and Garber 2014). This dataset demonstrates that these decisions are crucial. Lower nutritive value diets can lead to higher mortality rates, lower pregnancy rates, production of fewer offspring and a higher risk of predation (Proffitt et al. 2016). An analysis of 77 mammalian herbivores showed that larger animals better tolerate diets of lower nutritive quality because they can consume a greater volume of vegetation without increasing the efficiency of digestion (Müller et al. 2013). Larger herbivores also process their food more slowly, and are generally ruminants, whereas smaller hindgut fermenters feed selectively on the most digestible plants (Illius and Gordon 1992; Clauss et al. 2003). These data suggest that, across arid and equatorial regions, larger ruminant herbivores may be favoured by the lower nutritive values of the forage plants which grow there, whereas smaller hindgut fermenters may be favoured in temperate and tundra regions. There are other factors which also play important roles, including predation or poaching risk, competition, temperature stress and drought frequency (Gaston and Blackburn 1995; Cardillo and Bromham 2001).

Regional and inter-annual variability in climate generates corresponding variation in forage nutritive values (Grant et al. 2014; Ray et al. 2015). This variability influences animal migrations, for example wildebeest and zebra travel larger distances and remain within grazing patches for shorter period when forage is of high nutritive value (Hopcraft et al. 2014). Herbivores that do not migrate display the opposite pattern, since they spend more time in the same patch consuming the more nutritious forage plants (Laca et al. 1994). The spatial and temporal variation in forage plants shown here may contribute to explanations of optimal herbivore migration strategies and foraging behaviour. In addition, reduction in forage quality driven by climate change have been projected (Lee et al. 2017). Lower nutritive values in warmer bioclimatic zones adds further evidence to these projections and also suggests that future changes to forage nutritive values may modify migration and grazing strategies (Walther et al. 2002). Enteric methane production is also increased when ruminants consume lower quality forage, and methane emissions may also be influenced by these spatial and temporal patterns in forage nutritive values (Knapp et al. 2014).

Grazing lands have expanded to supply the growing demand for meat and dairy products, particularly across Asia and South America, and now cover 35 million km^2^ of the Earth’s surface (FAOSTAT, 2016). The majority of the world’s livestock are subject to permanent or seasonal nutritional stress (Bruinsma 2003). Poor animal nutrition impairs livestock productivity across many smallholder farms, particularly in Africa and the developing world (Thornton et al. 2011). It has been suggested that plantation crops and industrial by-products may enhance animal nutrition (Thornton and Herrero 2010; Herrero et al. 2013). However, this dataset demonstrates that assessments of the nutritive values of forage plants may identify species with useful nutritive traits. This analysis was not limited to the developing world, and this database summarising the nutritive values of forage plants, may be used to identify species which can be cultivated across different regions according to the nutritive values needed. In the USA, for example, the nutritive values of forage plants has declined over the past 22 years and this decline has been linked with drought, rising atmospheric CO_2_ concentrations, and sustained nutrient export (Craine et al. 2017). Forage species of high nutritive value which grow in warmer and drier regions could be selected. Future responses to global changes must also be considered, with warming, modified rainfall patterns, fertilisation and CO_2_ enrichment associated with changes to forage plant productivity and nutritive quality (Milchunas et al. 2005; Craine et al. 2010; Lee et al. 2010, 2014).

### Grasses

Grasses grow rapidly and are frequently described as the most tolerant group to herbivory (Wang et al. 2012, 2013). In the year 2000, 48 % (2.3 billion tons) of the biomass consumed by livestock was grass, followed by grains (1.3 billion tons). The remainder of livestock feed (0.1 billion tons) was derived from the leaves and stalks of field crops, such as corn, sorghum and soybean (Herrero et al. 2013). Grasses were the most variable group in this dataset. This was, in part, because grasses comprised the majority of the data points. However, these data revealed the extent by which grasses may vary in their nutritive values in terms of DM (11-97%), water (3-89%), protein (2-36%), fibre (defined by NDF; 29-90%), minerals (defined by ash; 2-19%) and lignin contents (1-21%).

It has been shown that birds, amphibians, reptiles, mammals and arthropods select grasses based on nutritive values (Simpson and Raubenheimer 1993; Simpson et al. 2004). In a study of wild grass-consuming herbivores across Africa, diet composition was shown to be consistent within consumer species but varied between consumer species, whilst total biomass intakes were constant indicating that grass nutritive characteristics were important determinants of herbivore body sizes (Kartzinel et al. 2015). Such variation may contribute to niche segregation and to the coexistence of large herbivores of relatively similar body mass, as observed in mountain ecosystems (Redjadj et al. 2014). This dataset provides further evidence for forage driven niche segregation among herbivores by quantifying the substantial capacity for variation in nutritive values between forage species and functional groups, as assessed by different nutritive metrics.

### Legumes

Cultivating herbaceous legumes has been proposed as a method for improving the protein content of pasture, particularly in the arid and equatorial rangelands of Asia, Africa and Latin America (Derner et al. 2017). Herbaceous legumes are planted increasingly frequently across temperate and tundra regions, in part because of their elevated protein content, improving meat and milk protein, and in part because of enhanced soil nitrogen availability, reduced fertiliser usage and reduced nitrous oxide emissions (Lüscher et al. 2014). Biomass production can be increased by fertilisation and legumes can be tolerant to increased salinity, albeit at low concentrations (Zouhaier et al. 2016). Some wild herbivores are specialist legumes feeders and have different nutritive requirements from generalists or those which consume plants in other functional groups (Karowe 2007). This dataset demonstrates that legumes generally provide greater concentrations of protein, supporting their use as a component of livestock fodder. The magnitude of the increased protein content of legumes was greatest across arid and equatorial regions, where the benefits of additional protein in human diets may be greatest (Tilman and Clark 2014).

Legumes also generally contain lower levels of fibre and higher concentrations of minerals than grasses. This may be driven by the branched venation patterns of the leaves of herbaceous legumes compared with the parallel system of vascular bundles running the length of grass leaves combined with their shorter habit which requires less structural fibre (Jung and Allen 1995). Herbaceous legumes may therefore have the combined nutritive benefits across arid and equatorial regions of greater protein and lower fibre contents compared with grasses. It should be noted that some grasses contained high protein, high minerals, low fibre and low lignin contents and there was no difference in mean digestibility between grasses and legumes. Care must be taken to consider the full suite of nutritive metrics, including their positive or negative effects on overall plant productivity, when selecting herbaceous legumes for use as livestock fodder (Wagner et al. 2016).

### Tree foliage

Trees and shrubs can deliver forage alongside several other ecosystem services, including carbon storage, soil fertility, flood defence and biodiversity enhancement, and there has been recent research interest in quantifying the benefits of silvopastoral livestock systems (Santos et al. 2016), particularly in restoring degraded pasture (Yamamoto et al. 2007). Trees can provide supplementary forage, because tree foliage has different nutritional profiles to other functional groups and trees are also productive during the times of the year when other plants are scarce (Salem et al. 2006). Tree leaves are also important foods for arboreal wild herbivores, such as primates, rodents, and marsupials, which often select foliage of high nutritive value and avoid leaves with high tannin or lignin contents (Farmer, 2014). Across this dataset, tree foliage was generally higher in terms of lignin and fibre contents than the other functional groups, and the ranges of values of DM (22-93%), water (7-78%), protein (10-25%), fibre (as defined by NDF; 33-68%), minerals (4-22%) and lignin contents (3-21%), were generally lower than the grasses and in line with those found for herbaceous legumes. High lignin and fibre contents of tree foliage could limit livestock productivity, however, it has been shown that cattle consuming tree foliage as a supplement to grass can continue to deliver high milk and meat yields (Andrade et al, 2008). Some tree species can regrow foliage following herbivory, however, increased light intensity can increase tannin concentrations (Nabeshima et al. 2003). As with legumes, care must be taken in selecting tree species for inclusion in cattle diets, in particular by quantifying lignin and tannin contents. Understanding the roles of different nutritive components may also provide a deeper understanding of arboreal herbivore population dynamics and behaviour (Coley and Barone 1996).

### Herbaceous non-legumes

Generally the productivity of non-leguminous herbaceous plants is much lower than the other functional groups, limiting their use for livestock fodder (Kallah et al. 2000; Elgersma et al. 2014). However. the advantages of cultivating herbaceous non-legumes include the prevention of weed establishment, the enhancement of conservation value, the extension of grazing periods and elevated forage mineral contents (Pirhofer-Walzl et al. 2011). There were few nutritive differences between the herbaceous non-legumes and the other functional groups, although herbs generally contained less fibre than grasses, less protein than legumes and less lignin and fibre than trees. Planting some herbaceous species can enhance livestock productivity and can also modify the taste of dairy products (Vasta et al. 2008). Many wild herbivores also utilise herbaceous plants for food, particularly arthropods (Siemann et al. 1999). This dataset highlights that some herbaceous plants may offer nutritional costs and benefits to livestock and wild herbivores, and studying their nutritive values may provide insight into herbivore population dynamics. However, due to the low representation of this group in the dataset, further work is required to fully quantify variation in the nutritive value of herbaceous non-legumes (Gasson and Cutler 1990).

## Conclusions

This dataset reveals the extent by which different species of forage plants can vary in their nutritive value to herbivores. Some forage plant species were highly nutritious containing high concentrations of protein and minerals and low concentrations of fibre and lignin, resulting in high digestibility values. This highlights the importance of foraging decisions made by wild and domesticated herbivores. This dataset also demonstrates the capacity for improved livestock forage if species selection is based on forage quality. This may also be important for conservation efforts, if the nutritional requirements of the target organisms are well understood. Multiple agronomic nutritive metrics were considered in this analysis, and many were auto-correlated, but fibre content was the best predictor of low quality forage, as defined by low digestibility values. High fibre content or low digestibility may be the best proxy for poor quality forage. Forage quality was also lower in warmer and drier arid and equatorial regions suggesting that the availability of high quality forage across these regions is low. This information may contribute to explanations of variation in optimal herbivore body sizes, migration behaviour and grazing patterns. Projections of the effects of climate change on plant/herbivore interactions should consider future changes to forage plant nutritive values and plant species composition.

## Acknowledgements

Thanks to Aaron Davis for advice on preparation of this manuscript.

